# Evolution of *Shewanella oneidensis* MR-1 in competition with *Citrobacter freundii*

**DOI:** 10.1101/2023.06.15.545074

**Authors:** Biyi Zhao, Geng Chen, Wei Chen, Yong Xiao

## Abstract

Interspecific competition is one of the most important metabolic interactions within microbial communities, and exoelectrogenic bacteria that can conduct extracellular electron transfer act essential roles in nature and engineering systems for pollutants removal. In the present study we investigated the long-term impact of substrate competition from non-exoelectrogenic *Citrobacter freundii* An1 on exoelectrogenic *Shewanella oneidensis* MR-1. Without additional electron acceptor or with electron acceptor of oxygen, *C. freundii* An1 typically suppressed the growth of *S. oneidensis* MR-1. In contrast, *S. oneidensis* MR-1 grown better with electron acceptor of ferrihydrite by taking advantage of extracellular electron transfer. However, the presence of ferrihydrite did not enhance the ferrihydrite reduction of *S. oneidensis* MR-1 after the 160 d-acclimation. The whole genome resequencing showed a complex evolution of *S. oneidensis* MR-1 when the strain faced the competition from *C. freundii* An1 for substrate.

## 1 INTRODUCTION

Extracellular electron transfer (EET), the main mode of extracellular metabolism in microorganisms, is important for geochemical cycles because of the process that converts solid metal minerals (Hernandez and Newman 2001) and has a wide range of applications in the environmental field (Nazari et al. 2020). Electron acceptors also drive the evolution of organisms on Earth, and electron acceptors in the environment influence the respiratory metabolism of microorganisms and determine their ecological niche distribution. At the same time, interspecific competition is one of the most prevalent interactions affecting community structure in communities (Machado et al. 2021). It has been shown that substrate competition is an important driver of microbial evolution and influences changes in microbial ecological niches. However, the response of microbes to different electron acceptors in a substrate competitive environment and how electron acceptors affect microbial evolution is not yet understood.

In a previous study, we have showed that competition for substrate enhanced the EET of *Shewanella oneidensis* MR-1. In the present study, we constructed the synthetic consortium between *S. oneidensis* MR-1 (a strains capable of respiration using a wide range of electron acceptors) and *Citrobacter freundii* An1, using lactate as a substrate, to investigate the mechanisms of microbial evolution in different electron acceptor systems under long-term substrate competition stress.

## 2 MATERIALS AND METHODS

### 2.1 Bacterial strains and culture conditions

The two strains are *S. oneidensis* MR-1 and *C. freundii* An1 that we have used in a previous study (Xiao et al. 2021). Luria-Bertani (LB) broth (pH 7.0) was use for strains recovery from -80 °C. This medium consists of tryptone (10 g/L), yeast extract (5 g/L), and NaCl (10 g/L). A starter culture was derived from a single colony on a freshly streaked LB plate which was made from LB broth with another 1.5% agar. The single colony was then transferred into 100 mL of LB broth and incubated for 16 h. This culture was regarded as the original population (designated as 0d-MR-1 and 0d-An1).

For the long-term acclimation experiments, another medium was prepared as following: 5% LB, 50 mM Na_2_HPO_4_, 5 mM sodium lactate, pH 7.0. The culture temperature was set at 30 °C, and the flasks was shaken at a speed of 150 rpm. Ferrihydrite was synthesized following a previous study (Lovley Derek and Phillips Elizabeth 1986).

### 2.2 Long-term acclimation experiment

According to the difference of electron acceptors, three groups were set for the long-term acclimation experiments. All groups with different treatments were shown in Table 1, and each treatment has three replicates. For the mixed culture with both *S. oneidensis* MR-1 and *C. freundii* An1, bacterial cells of about 40% was *S. oneidensis* MR-1 based on the result from blood corpuscle counting. During the long-term acclimation, 1% of parent cells was used to inoculate the fresh medium. Due to different growth in each group, the subculturing cycles of anaerobic, oxygen, and ferrihydrite group were about 7, 4, and 10 d, respectively. Samples collected at 45, 80, and 160 d were transferred into fresh LB broth with 25% glycerol and stored at -80 °C. *S. oneidensis* MR-1 and *C. freundii* An1 in mixed culture were then isolated from LB plate and identified by 16S rRNA gene sequencing, and each sample had at least 3 replicated colonies.

**Table 1.** Experiment groups and treatments in the long-term acclimation Electron.

| Group | Electron acceptor | Strains | Treatment name |
| --- | --- | --- | --- |
| Anaerobic control | none | *S. oneidensis* MR-1 | MR-1-Anaerobic control |
|  |  | *C. freundii* An1 | An1-Anaerobic control |
|  |  | *S. oneidensis* MR-1 and *C. freundii* An1 | Mix-Anaerobic control |
| Oxygen | oxygen | *S. oneidensis* MR-1 | MR-1-Oxygen |
|  |  | *C. freundii* An1 | An1-Oxygen |
|  |  | *S. oneidensis* MR-1 and *C. freundii* An1 | Mix-Oxygen |
| Ferrihydrite | ferrihydrite | *S. oneidensis* MR-1 | MR-1-Ferrihydrite |
|  |  | *C. freundii* An1 | An1-Ferrihydrite |
|  |  | *S. oneidensis* MR-1 and *C. freundii* An1 | Mix-Ferrihydrite |

### 2.3 Analysis of bacterial communities

Mixed culture samples collected at 0, 45, and 160 d from three groups were centrifuged at 12000 g for 3 min to collect cells. The 16S rRNA gene sequencing was performed on Illumina 300 platform by Biomarker Technologies Corporation (Beijing, China).

### 2.4 Whole genome resequencing and bioinformatics analysis

A total of 38 colonies were randomly selected for whole genome resequencing on Illumina 300 platform by Biomarker Technologies Corporation (Beijing, China). Two colonies from 0d-MR-1 and 0d-An1, and the others from the 9 treatments at 160 d. The bioinformatics analysis was performed following our previous study (Xiao et al. 2020).

## 3 RESULTS AND DISCUSSION

### 3.1 Impact of electron acceptors on bacterial communities

To investigate how electron acceptor impacts the community composition changes in the Mix groups, the relative abundance of *S. oneidensis* MR-1 in the Mix groups at 0, 45, and 160 d was examined using 16S rRNA gene-based amplicon high-throughput sequencing (Figure 1A). In the 0 d-Mix group, i.e., the initial inoculum percentage of each culture system, *S. oneidensis* MR-1 had an abundance of 38.1%. After 45 d of incubation, the relative abundance of *S. oneidensis* MR-1 in all Mix-groups decreased: 7.8±1.9% in the Mix-Anaerobic control group without the applied electron acceptor, 15.0±1.5% in the Mix-Oxygen group, and 29.9 ± 4.1% in the Mix-Ferrihydrite group. Then, the relative abundance of *S. oneidensis* MR-1 in Mix-Anaerobic control and Mix-Oxygen group further decreased to at 160 d 5.1±1.0% and 11.9±0.9%, respectively. In contrast, its abundance in the Mix-Ferrihydrite group showed an increase to 35.6±10.6% from 45 d to 160 d. Since *C. freundii* An1 is not able to conduct extracellular electron transfer (EET), it can not use ferrihydrite as electron acceptor to support its growth as that *S. oneidensis* MR-1 does. It can be inferred that, during the intense competition with *C. freundii* An1, *S. oneidensis* MR-1 uses ferrihydrite as an electron acceptor for EET, which is beneficial to its growth.

**Figure 1.**
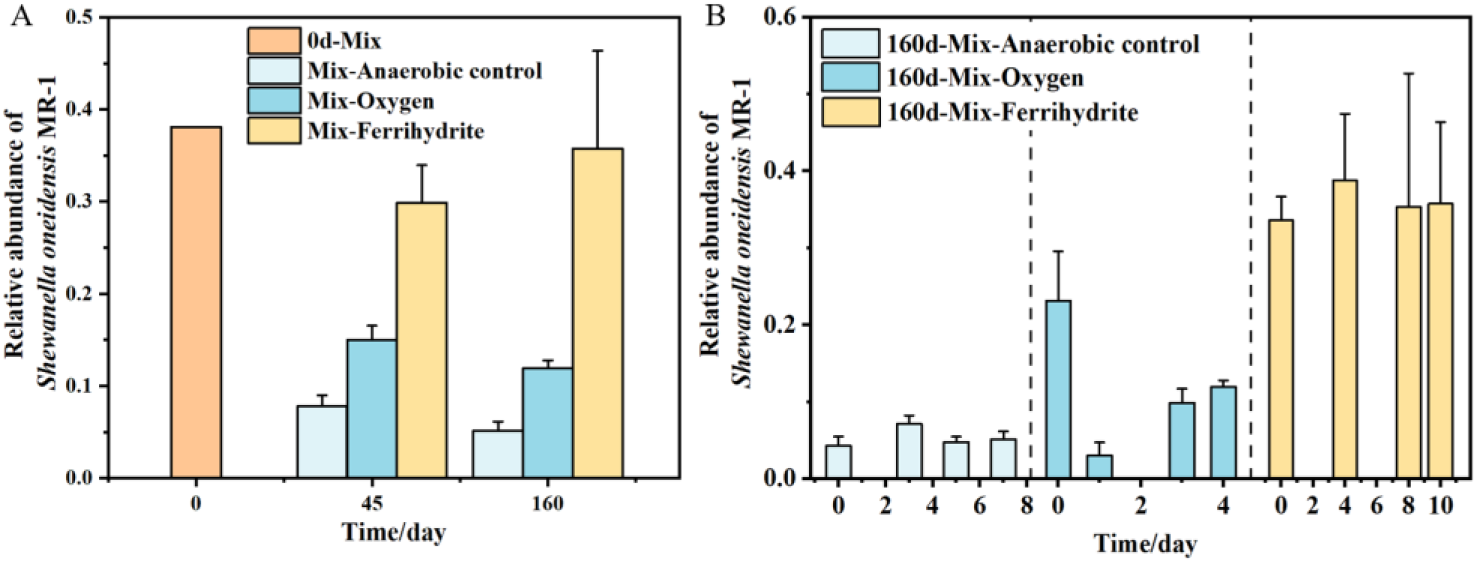
The relative abundance of *S. oneidensis* MR-1 in the Mix groups. A, changes during the long-term acclimation; B, changes in a single cycle at 160 d.

To investigate the competition process within a single cycle, the changes of relative abundance of *S. oneidensis* MR-1 in the Mix groups were further determined at around 160 d (Figure 1B). It was found that the relative abundance of *S. oneidensis* MR-1 in Mix-Anaerobic control group and Mix-Ferrihydrite group fluctuated less during this single cycle. The relative abundance of *S. oneidensis* MR-1 was maintained at about 5% in Mix-Anaerobic control group, while it could be maintained at around 35% in Mix-Ferrihydrite group. In contrast, the abundance of *S. oneidensis* MR-1 in 160d-Mix-Oxygen group was lower than that in Mix-Ferrihydrite group and showed a big fluctuation during the whole cycle. This result further suggested that for *S. oneidensis* MR-1, ferrihydrite is beneficial to its growth while competing with *C. freundii* An1 for substrate.

### 3.3 Competitive strategy of *S. oneidensis* MR-1 against oxygen

To investigate the competitive strategy of *S. oneidensis* MR-1 in the presence of oxygen, the growth curves of the ancestral *S. oneidensis* MR-1 (i.e. 0d-MR-1) and the 160 d-evolved *S. oneidensis* MR-1 isolated from groups MR-1-Oxygen and Mix-Oxygen were measured (Figure 2A). The results showed that there was no significant change in the growth rate of the 160d-Oxygen group compared with 0d-MR-1. However, the three isolates from 160d-Mix-Oxygen group showed different growth trends: 160d-Mix-Oxygen-MR-1_1 had a faster growth rate at the beginning of the culture and also entered the decay phase more quickly at the later stage; 160d-Mix-Oxygen-MR-1_2 could reach a maximum OD_600_ only 0.093, which is about one-fifth of the maximum OD_600_ of the 0d-MR-1; while for the 160d-Mix-Oxygen-MR-1_3 group, its maximum OD_600_ is 0.437, which is 20% less than that of 0d-MR-1. It can be inferred that after 160 d-acclimation, *S. oneidensis* MR-1 developed different strategies to compete with *C. freundii* An1 which grows better at the presence of oxygen.

**Figure 2.**
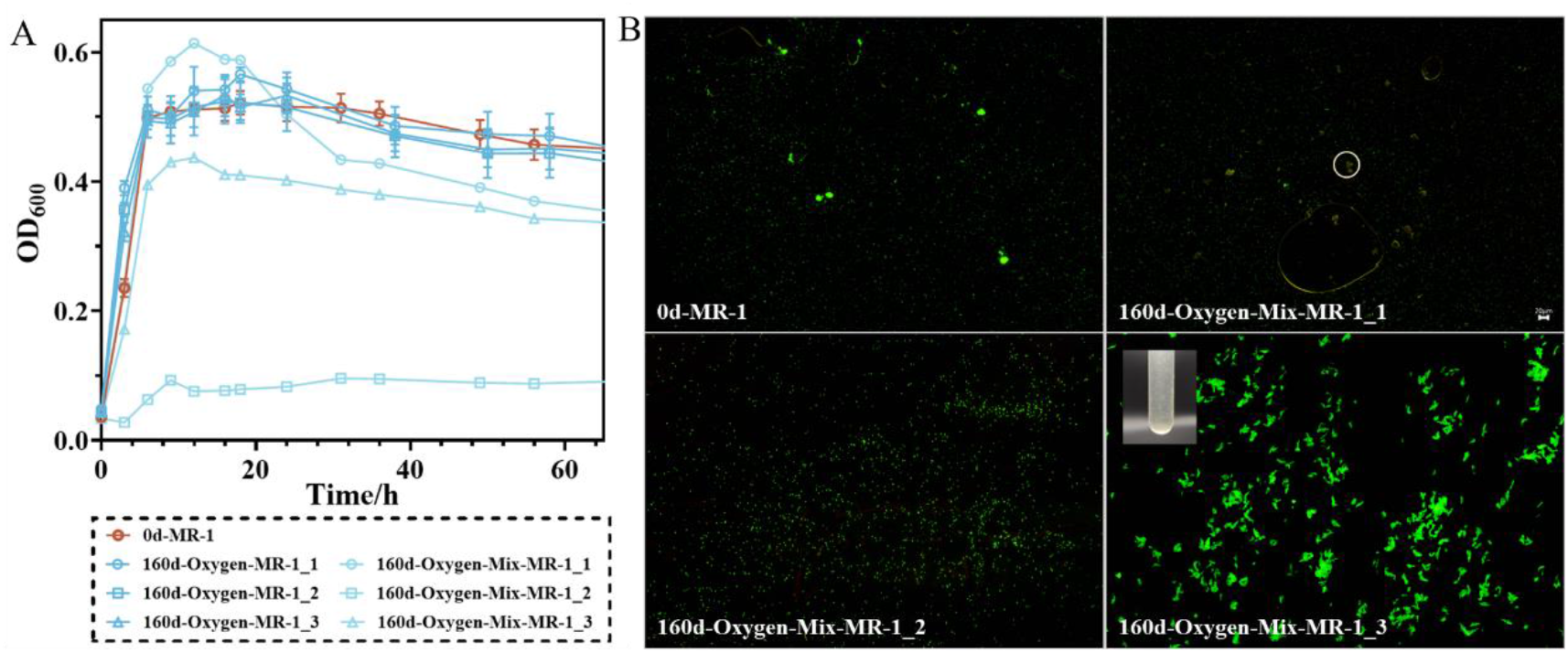
Growth curve (A) and morphology (A) of *S. oneidensis* MR-1 in groups using oxygen as electron acceptor.

Microbial aerobic respiration using oxygen as an electron acceptor produces large amounts of superoxide and hydrogen peroxide byproducts which can cause damage to cells (Imlay 2013). It has been shown that growth under aerobic condition triggers self-aggregation of *S. oneidensis* MR-1 to reduce oxidative stress and promote anoxic condition within the aggregates for anaerobic respiration and thus protecting the cells (McLean et al. 2008). Meantime, it has been shown that co-culture of microorganisms can promote the coexistence of species through spatial heterogeneity and local adaptation, i.e., the separation of ecological niches can promote the survival of each species in the co-culture system to prevent the extinction phenomenon (Lankau 2011). The formation of agglomerates separates the ecological niches of different microorganisms. Therefore, it is hypothesized whether the formation of agglomerates is a strategy to resist oxygen stress and to facilitate the survival of *S. oneidensis* MR-1 in a competitive environment by promoting the separation of ecological niches.

To verify the above conjecture, we stained the cells of 0d-MR-1 and *S. oneidensis* MR-1 from 160d-Mix-Oxygen group after 24 h of culture using a fluorescence microscopy imaging (Figure 2B). There were few and small aggregates in 0d-MR-1, and most of them were spherical aggregates with a maximum diameter of about 20 μm. For isolate of 160d-Mix-Oxygen-MR-1_1, it had only very few agglomerates whose size were mostly 20 μm. For isolate of 160d-Mix-Oxygen-MR-1_2, the cells were basically dispersed and did not appear to be aggregated. For the isolate of 160d-Mix-Oxygen-MR-1_3, there were obvious cell clusters with two morphologies: flocculent and (ellipsoidal) spherical, with flocculent sides up to 100 μm in length and spherical shapes up to 500 μm in diameter. These results indicated that *S. oneidensis* MR-1 developed different strategies against the stress from oxygen and the competition from *C. freundii* An1.

### 3.3 EET capacity change of *S. oneidensis* MR-1

Microorganisms can use different electron acceptors for respiratory metabolism in different modes. When using ferrihydrite as electron acceptor, *S. oneidensis* MR-1 can get some advantage to compete with *C. freundii* An1. We therefore investigated the EET capacity of *S. oneidensis* MR-1 after 160 d co-cultivation with *C. freundii* An1 as indicated by ferrihydrite reduction (Figure 3). It seemed that long-term acclimation with ferrihydrite did not necessarily enhance the EET capacity, as well as oxygen and the condition without any electron acceptor.

**Figure 3.**
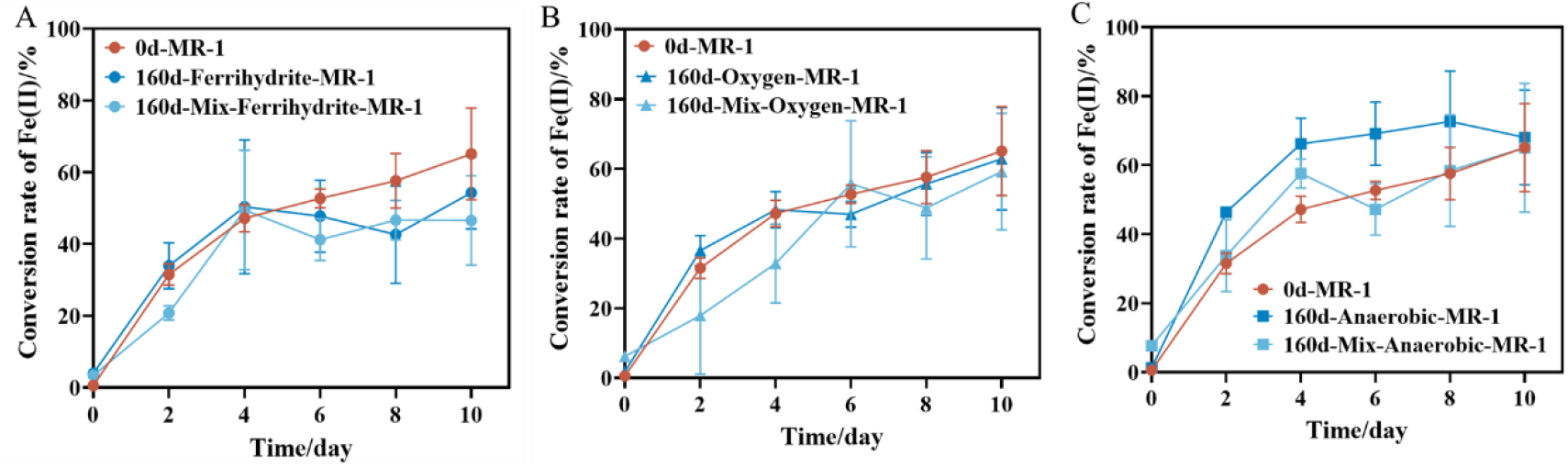
EET capacity change of *S. oneidensis* MR-1 after 160 d acclimation using ferrihydrite (A), oxygen (B), and none (C) as electron acceptors, respectively.

### 3.4 Resequencing illustrates the survival mechanism of *S. oneidensis* MR-1

After being acclimated for 160 d, *S. oneidensis* MR-1 in pure and co-culture groups were used for whole-genome resequencing to evaluate its adaptive mutation against different electron acceptors. More nonsynonymous mutations was found in the co-culture groups than that in the pure groups (Figure 4A). This result suggests that the substrate competition with *C. freundii* An1 may have caused some stress to *S. oneidensis* MR-1 and sequentially promoted more nonsynonymous mutations in *S. oneidensis* MR-1. Functional enrichment (GO enrichment) analysis (Figure 4B) and KEGG metabolic pathway enrichment analysis (Figure 4C) were performed on the nonsynonymous mutant genes of *S. oneidensis* MR-1 in the Mix-Oxygen group. The mutations were found to be mainly associated with the cellular chemotaxis process and regulatation of the synthesis (arginine) and metabolism (alanine, aspartate, glutamate and tryptophan) of amino acids related to cellular respiration, and the others mainly related to ribosomal synthesis process.

**Figure 4.**
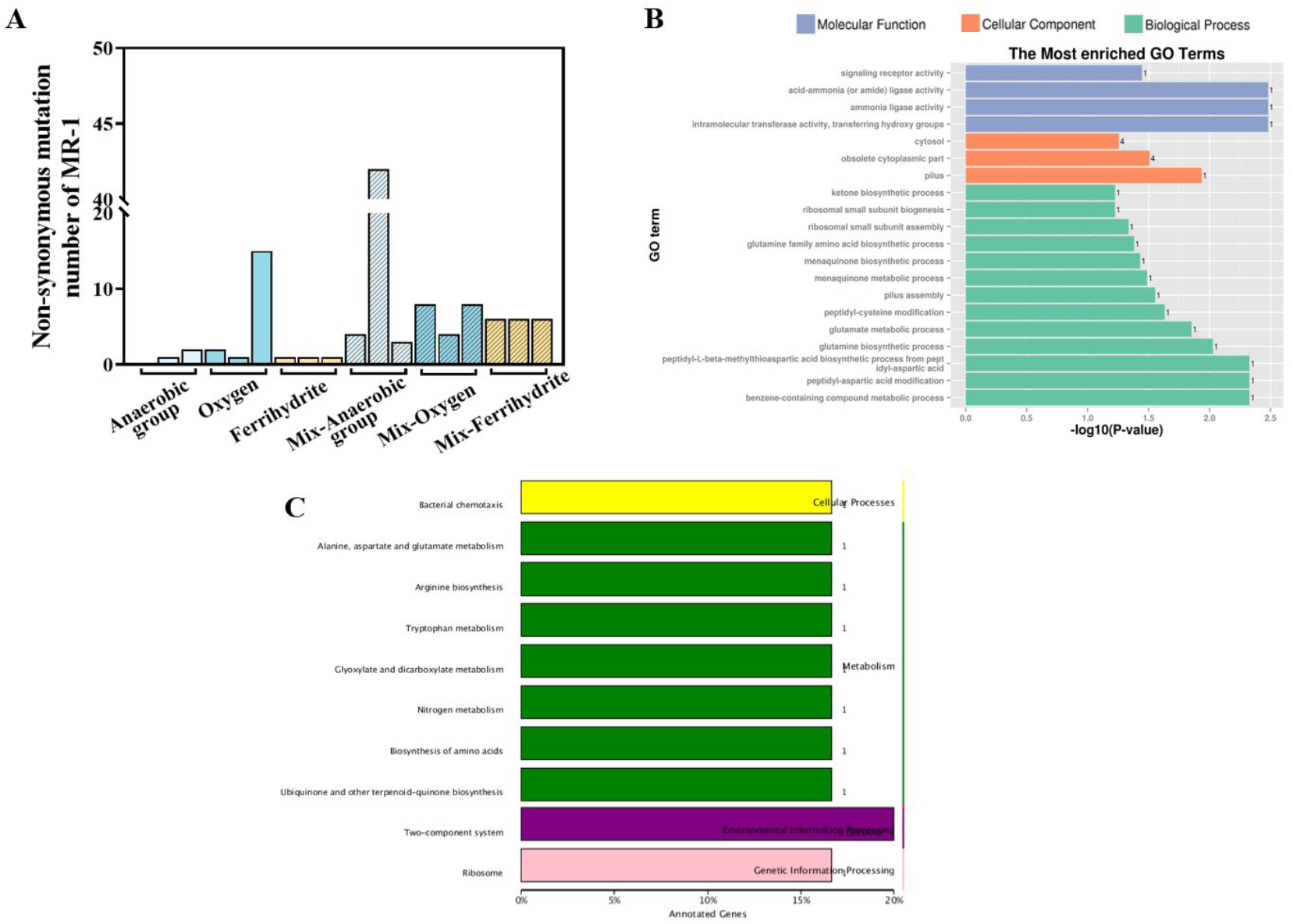
Whole-genome resequencing of *S. oneidensis* MR-1 find adaptive mutation against different electron acceptors. A, SNP number of *S. oneidensis* MR-1 in pure culture or co-culture from different treatments; B, GO enrichment analysis of *S. oneidensis* MR-1 from co-culture using oxygen; C, KEGG analysis of *S. oneidensis* MR-1 from co-culture using oxygen.

We further analyzed the nonsynonymous mutation sites of *S. oneidensis* MR-1 isolated after long-term (160 d) substrate competition in different electron acceptor groups. In systems where oxygen is the electron acceptor, mutations involving resistance to oxygen toxicity occur in both pure and co-culture systems because oxygen is toxic to microorganisms (Table 2). Gene *rfbB* is associated with the synthesis of biofilm polysaccharides (McLean et al. 2008). Gene *pilY* encodes a Type IV ciliate systemic adhesion protein which is associated with microbial motility and affects biofilm formation (Persat 2017, Waite et al. 2005). Gene *mgtE* encodes a magnesium ion transport protein that is associated with microbial anaerobic respiration (Merino et al. 2001) and affects microbial adhesion (Merino et al. 2001). Gene *menH* is related to the biosynthesis of methylnaphthoquinone (Sepúlved a-Cisternas et al. 2018, Wan et al. 2017). Gene *SO_3642* is associated with microbial tropism and may also affect biofilm formation in *S. oneidensis* MR-1, while *SO_1414* encodes flavin flavocytochrome c, and both menaquinone and cytochrome *c* are involved in the electron transport chain and are associated with the anaerobic respiration process of *S. oneidensis* MR-1 (Alonso et al. 2014, Nowicka and Kruk 2010). *dnrN* encodes an iron-sulfur cluster repair protein which may mitigate oxidative stress processes and repair oxidative damage in microbes (Overton et al. 2008). *SO_4413* encodes kynureninase which is associated with the synthesis of the antioxidant kynurenine in microorganisms and may inhibit the production of reactive oxygen species by promoting the synthesis of kynurenine (Genestet et al. 2014).

**Table 2.**
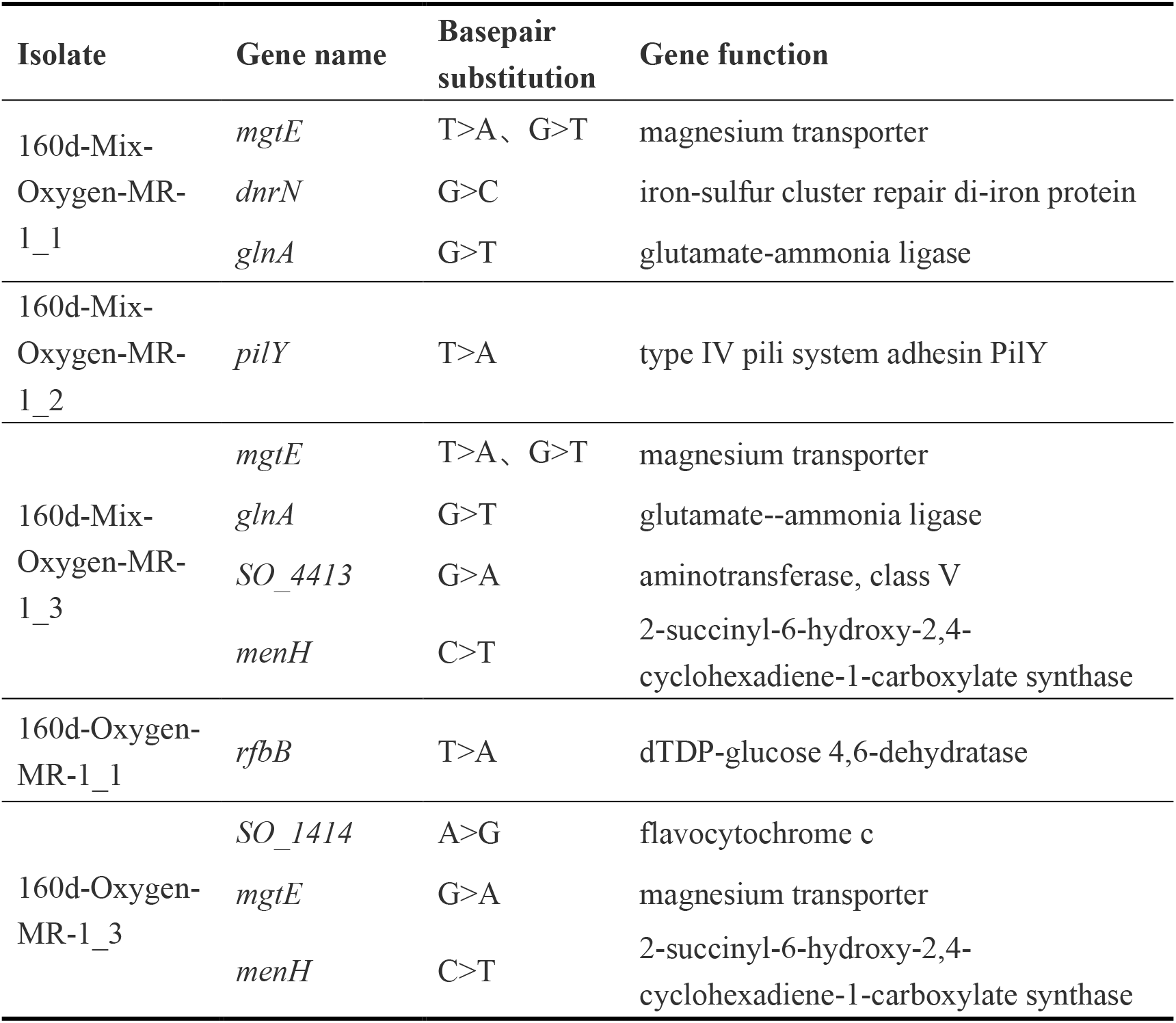
Nonsynonymous mutant sites after 160-day cultivation in oxygen group.

In summary, *S. oneidensis* MR-1 developed several strategies of promoting biofilm formation, cellular self-aggregation, anaerobic respiration and metabolic activity to survive when oxygen was used as electron acceptor, and substrate competition with *C. freundii* An1 further promoted more abundant adaptive mutations in *S. oneidensis* MR-1. In contrast, the nonsynonymous mutations seemed to be random in *S. oneidensis* MR-1 when the electron acceptors are ferrihydrite and none. This may be related to smaller competitive stress on *S. oneidensis* MR-1 in these two groups since *C. freundii* An1 did not grow well in these two groups.

## ACKNOWLEDGMENTS

This study was supported by grants from the Youth Innovation Foundation of Xiamen (3502Z20206089), National Natural Science Foundation of China (22276183), and the Natural Science Foundation of Fujian for Distinguished Young Scholars (2021J06036).

## Notes

### Competing Interest Statement

The authors have declared no competing interest.

### Summary of Updates

layout update

